# Mechanisms of protein kinase C epsilon down-regulation by transforming growth factor-beta in lung cancer cells

**DOI:** 10.1101/2020.10.28.359588

**Authors:** Victoria Casado-Medrano, Martin J. Baker, Mariana Cooke, Marcelo G. Kazanietz

**Affiliations:** Department of Systems Pharmacology and Translational Therapeutics, Perelman School of Medicine, University of Pennsylvania, Philadelphia, PA 19104, USA; Department of Medicine, Einstein Medical Center Philadelphia, Philadelphia, PA 19141, USA

**Keywords:** PKCε, TGF-β, proteasome, autophagy, lysosome, ubiquitination

## Abstract

Protein kinase C epsilon (PKCε), a diacylglycerol (DAG)/phorbol ester-regulated PKC isoform, has been widely linked to oncogenesis and metastasis. PKCε plays important roles in the regulation of motility and invasiveness in non-small cell lung cancer (NSCLC). We previously reported that this kinase becomes prominently down-regulated upon TGF-β-induced epithelial-to-mesenchymal transition (EMT), which leads to prominent phenotypic changes. While the phorbol ester PMA causes down-regulation of PKCα, δ and ε within hours, TGF-β requires at least 4 days to reduce the expression levels of PKCε without affecting the expression of other PKCs, an effect that parallels the acquisition of a mesenchymal phenotype. Despite the prominent transcriptional component involved in EMT, we found that PKCε down-regulation does not involve changes in PKCε mRNA levels and was entirely independent of transcriptional activation of the *PRKCE* gene. Further mechanistic analysis revealed that the reduction in PKCε expression is dependent on proteasomal and endolysosomal pathways, but independent of autophagy processing mechanisms. Site-directed mutagenesis of Lys312 and Lys321 in PKCε prevented its down-regulation in response to either TGF-β or the phorbol ester PMA. The shift in PKCε isozyme levels depending on cell plasticity underscores relevant functional consequences by modulating the expression of this oncogenic/metastatic kinase and highlights key roles of protein stability mechanisms in the control of PKCε phenotypic outcomes.

## INTRODUCTION

The protein kinase C (PKC) family of serine/threonine kinases are the best characterized cellular effectors for the lipid second messenger diacylglycerol (DAG) and the phorbol ester tumor promoters. These kinases play fundamental roles in normal cell physiology and have been widely implicated in disease, including cancer. Based on their biochemical and structural properties, the 10 members of the PKC family have been classified into “conventional/classic” or cPKCs (α, βI, βII and γ), “novel” or nPKCs (δ, ε, θ and η) and atypical or aPKCs (ι/λ and ζ). Only cPKCs are responsive to calcium, whereas both cPKCs and nPKCs are activated by DAG and phorbol esters. The accepted general paradigm of PKC activation is that DAG generated upon stimulation of membrane receptors promotes cPKC and nPKC translocation to the plasma membrane, inducing a conformational change that leads to phosphorylation of specific PKC substrates (1–4). DAG is a short-lived second messenger, thus receptor-mediated translocation of PKCs is a rapid and transient event. On the other hand, lipophilic phorbol esters such as PMA (phorbol 12-myristate 13-acetate) or synthetic DAG analogues cause sustained PKC translocation to membranes. The lasting association of PKCs with membranes leads to their slow down-regulation in expression. Individual PKC isozymes follow distinctive patterns of translocation and down-regulation depending on the ligand and cell type (5–9).

While acute stimulation of PKC with phorbol ester and related analogues is widely used to recapitulate DAG-mediated responses in cellular models, in many cases PKC-mediated functions are examined in response to long-term treatment with phorbol esters, thus suggesting potential relationships with PKC loss of expression. A typical example is the widely used multistage skin carcinogenesis experimental paradigm that involves multiple weeks of topical PMA applications after a single carcinogen topical treatment (1, 10). Since individual PKC isozymes have opposing roles in tumorigenesis either as promoters or suppressors of growth, the controversy still remains as to whether phorbol ester-mediated tumor promotion depends on the persistent activation of a tumor promoting PKC or the down-regulation of a tumor-suppressing PKC (1, 11–13). Interestingly, loss of PKC isozyme expression also takes place in a number of physiological processes such as differentiation (14, 15), underscoring the potential relevance of PKC degradation in cell fate determination. Whereas PKC down-regulation by phorbol esters has been studied in multiple contexts, the detailed mechanisms involved in enzyme degradation remain partially understood. The potential utilization of multiple degradation pathways in PKC isozyme down-regulation adds new layers of complexity to the regulation of PKC activity.

Among the different member of the PKC family, PKCε has been identified as a prooncogenic kinase and a cancer biomarker (1, 16). This nPKC is distinctly up-regulated in various epithelial cancers, including breast, head and neck, prostate and lung cancer. Recent studies have pointed to key roles for PKCε in cell survival, mitogenesis, and motility, and in upstream cancer signaling pathways, namely NF-κB, Akt, Stat3 and Rac pathways (12, 16–24). Interestingly, tumor growth could be abrogated by genetic deletion or silencing of PKCε, as well as by treatment with PKCε inhibitors (25–29). Furthermore, emerging evidence suggests potential roles for PKCε in EMT. While PKCε is required for EMT in mammary cancer models, it is dispensable for transforming growth factor-β (TGF-β)-induced EMT in non-small cell lung cancer (NSCLC) cellular models (30, 31). In a recent study, we reported that PKCε becomes markedly down-regulated by NSCLC cells progressively transformed to a mesenchymal state, an effect that is not observed for other DAG/phorbol ester responsive PKCs expressed in NSCLC, *i.e.* PKCα and PKCδ. Loss of PKCε expression in mesenchymally transformed NSCLC cells leads to important phenotypic consequences, specifically changes in the activation status of Rho GTPases widely implicated in migration and invasiveness (31). Dissecting the mechanisms that control PKCε down-regulation may help elucidate important regulatory aspects for this oncogenic kinase.

In this study, we pursue a comprehensive analysis to demonstrate unique mechanisms responsible for PKCε down-regulation by TGF-β. Identifying the characteristic pathways for PKCε degradation offers opportunities to disentangle the intricacies of PKC regulation and function as well as potentially helping in the future development of tools to target this oncogenic kinase.

## MATERIAL AND METHODS

### Cell lines and reagents

All cell lines were obtained from ATCC (Manassas, VA) and are fully authenticated (see ATCC homepage). Human NSCLC (A549, H358), prostate cancer (PC3), colon cancer (HCT116) and ovarian cancer (SKVO3) cell lines were cultured in RPMI medium supplemented with 10% FBS, 100 U/ml penicillin, and 100 μg/ml streptomycin. The human pancreatic cancer cell line AsPC1 was cultured in DMEM medium supplemented with 10% FBS, 2 mM glutamine, 100 U/ml penicillin, and 100 μg/ml streptomycin. Human TGF-β was purchased from Peprotech (Rocky Hill, NJ). Cycloheximide, bafilomycin A1, chloroquine, bortezomib, dynasore, leupeptin, methyl-β-cyclodextrin, pepstatin A, E64, ALLN, MG-132 and puromycin were purchased from Sigma-Aldrich (St. Louis, MO). Geneticin was obtained from Gibco, Gaithersburg, MD).

### Western blot analysis

Western blots were carried out as previously described (32). The following antibodies were used: anti-PKCδ (catalog # 2058S), anti-PKCε (catalog # 2683S), anti-LC3B (catalog # 2775), anti-ATG5 (catalog # 12994), anti-ATG12 (catalog # 4180), anti-beclin (catalog # 4122), anti-mTOR (catalog # 2983), anti-caveolin (catalog # 3267), anti-clathrin (catalog # 4796), anti-HA (catalog # 3724S), anti-FLAG (catalog # 8146S), anti-HSC70 (catalog # 8444), anti-HSP90 (catalog #4877), anti-rabbit isotype control (catalog # 3900), anti-vimentin (catalog # 3390S, Cell Signaling Technology, Danvers, MA), anti-PKCα (catalog # sc-208, Santa Cruz Biotechnology, Dallas, TX), anti-vinculin (catalog # V9131, Sigma-Aldrich), anti-β-actin (catalog # A5441, BD Biosciences, Franklin Lakes, NJ) and anti E-cadherin (catalog # MAB1838, RD Systems, Minneapolis, MN).

### RNA interference (RNAi)

ON-TARGET Plus small interfering RNAs (siRNAs) were purchased from Dharmacon (Lafayette, CO). We used the following siRNAs: AGT5 (catalog #L-004374), ATG12 (catalog # L-010212), beclin (catalog # L-010552), HSC70 (catalog # J-017609), HSC90 (catalog #J-005186), caveolin (catalog #L-003467), and CLTC (catalog #L-004001). siRNA for mTOR (catalog #6381S) was obtained from Cell Signaling Technology (Danvers, MA). ON-TARGET Plus non-targeting pool (Catalog # D-001810) was used as a control. siRNAs were transfected with Lipofectamine RNAi/Max (Invitrogen-Life Technologies, Grand Island, NY), as previously described (33).

### Generation of PKCε mutants

Mutations in PKCε were introduced with the QuikChange XL site-directed mutagenesis kit (Stratagene, La Jolla, CA). The following primers were used for PCR: for 5’- CCGCTGTTGGTGATTCTGTCTGGGGTAACGC-3’ (forward) and 5’- GCGTTACCCCAGACAGAATCACCAACAGCGG-3’ (reverse) for K312R-PKCε; 5’- CCAGCAATGAGCTTTCTCCTTCTCTGGCCGC-3’ (forward) and 5’- GCGGCCAGAGAAGGAGAAAGCTCATTGCTGG-3’ (reverse) for or K321R-PKCε; Mutations were confirmed in all cases by DNA sequencing.

### Generation of stable cell lines

For stable depletion of PKCε, A549 cells were infected with PKCε shRNA Mission® lentiviral transduction particles (catalog # SHCLNM_005400) from Sigma-Aldrich according to the manufacturer’s protocol. Pools were selected with puromycin (1 μg/ml). We used the shRNA clone TRCN0000219726 that is designed against the 3’UTR. The target sequence is CTGCATGTTCAGGCATATTAT. As a control, we used a non-target control shRNA lentivirus (catalog # SHC00IV).

For the generation of stable A549 cell lines expressing various PKCs and mutants we used following plasmids: pcDNA3-PKCε-Flag (wild-type, Addgene, Cambridge, MA), pcDNA3-Flag-K312PKCε, pcDNA3-Flag-K321-PKCε, pcDNA3-Flag-K312/K321-PKCε. Plasmids were transfected using jetOPTIMUS as recommended by the manufacturer Polyplus (Illkirch, France) (Thermo Fisher Scientific, Waltham, MA). Selection of stable cell lines was carried out with geneticin. HA-ubiquitin pcDNA3 expression plasmid was purchased from Addgene.

### Luciferase assays

Cells in 12-well plates were transfected with 450 ng of a reporter plasmid for the PKCε gen (*PRKCE*) promoter (pLuc-PKCε, bp −1933 to bp +219 bp) (34). As control we used pLuc empty vector. The *Renilla* luciferase expression vector pRL-TK (50 ng, Promega, Madison WI) was co-transfected for normalization. Transfections were done with Lipofectamine 2000 (Invitrogen, Carlsbad, CA). After 24 h, the cells were serum starved for 24 h and treated with TGF-β (10 ng/ml) for 0-72 h or PMA (100 nM) for 0-16 h. Cells were then lysed with passive lysis buffer (Promega, Madison, WI) and luciferase activity was determined in cell extracts using the Dual-Luciferase reporter assay system (Promega, Madison, WI). Each experiment was performed in triplicate. The results were normalized to *Renilla* luciferase activity and expressed as relative luciferase units (RLU).

### Immunoprecipitation

A549 cell pellets were lysed with a lysis buffer containing 20 mM Tris-HCl pH 7.5, 150 mM NaCl, 1 mM EDTA, 1% Triton X-100, 1mM glycerophosphate, complete protease inhibitor cocktail (Roche, Basel, Switzerland, catalog # 4693116001) and phosSTOPTM inhibitor (Roche, catalog # 4906845001). For immunoprecipitation, cleared lysates were mixed with Protein G/A (50:50) recombinant protein G agarose beads (catalog #15920-010, Invitrogen, Carlsbad, CA) coupled with anti-HA (catalog #3724S, Cell signaling, Danvers, MA), anti-Flag-M2 (F3165, Sigma-Aldrich, St. Louis, MO) or IgG control. Samples were incubated overnight at 4 °C, and then beads were washed three tim*e*s in lysis buffer.

### Immunofluorescence

For immunofluorescence, cells growing on glass coverslides at low confluence (~20-30 %) were fixed with 4% formaldehyde and stained with the following antibodies: anti-PKCε (catalog # 2683S), anti-EEA1 (catalog #2411S), anti-Rab5 (catalog #3547T), or anti-Rab7 (catalog #9367T), As secondary antibodies we used Alexa 488 (catalog # A11001), Alexa 555 (catalog # A21428), and Alexa 549 (catalog # A-11042) (Thermo Fisher Scientific). The nuclei were counterstained with DAPI. Slides were visualized using the University of Pennsylvania Cell & Developmental Biology Microscopy Core Zeiss LSM 710 confocal microscope or Zeiss LSM880 confocal microscope with Airyscanner.

### Quantitative real-time PCR (Q-PCR)

Total RNA was extracted using the RNeasy kit (Qiagen, Valencia, CA). Reverse transcription of RNA was done using the TaqMan reverse transcription reagent kit (Applied Biosystems, Branchburg, NJ). Q-PCR amplifications were performed using an ABI PRISM 7300 Detection System in a total volume of 20 μl containing TaqMan Universal PCR Master Mix (Applied Biosystems), commercial target primers (300 nM), fluorescent probe (200 nM), and 1 μg of cDNA. PCR product formation was continuously monitored using the Sequence Detection System software version 1.7 (Applied Biosystems). The FAM signal was normalized to endogenous UBC (housekeeping gene).

### Statistical analysis

Statistical analysis was done with either *t*-test or ANOVA using GraphPad Prism 3.0. In all cases, a p-value <0.05 was considered statistically significant.

## RESULTS AND DISCUSSION

### TGF-β-induced PKCε down-regulation in NSCLC cells is independent of transcriptional/translational mechanisms

PKCε is highly expressed in human NSCLC and plays important roles in tumor growth and metastasis (21,24,25). We recently found that PKCε undergoes a profound down-regulation as NSCLC cells acquire a mesenchymal phenotype in response to TGF-β, an effect that leads to important phenotypic consequences (31). A time-course analysis of PKCε expression in A549 NSCLC revealed ~ 60% and 80% down-regulation after 4 and 6 days of TGF-β treatment (10 ng/ml), respectively. This effect is specific for PKCε, since the expression levels of the other DAG-responsive PKCs expressed in NSCLC cells, namely PKCα and PKCδ, remained unaffected after long-term TGF-β treatment (Fig. 1A). This isozyme specificity and kinetics of down-regulation is remarkably different from that induced by PMA (100 nM), which down-regulates all three DAG/phorbol ester-responsive PKCs a few hours after initiation of treatment. Indeed, down-regulation of PKCα, PKCδ and PKCε isozymes is readily detected 4 h after PMA treatment, reaching a maximum at 16-24 h (Fig. 1A), as previously reported (9).

**Figure 1.**
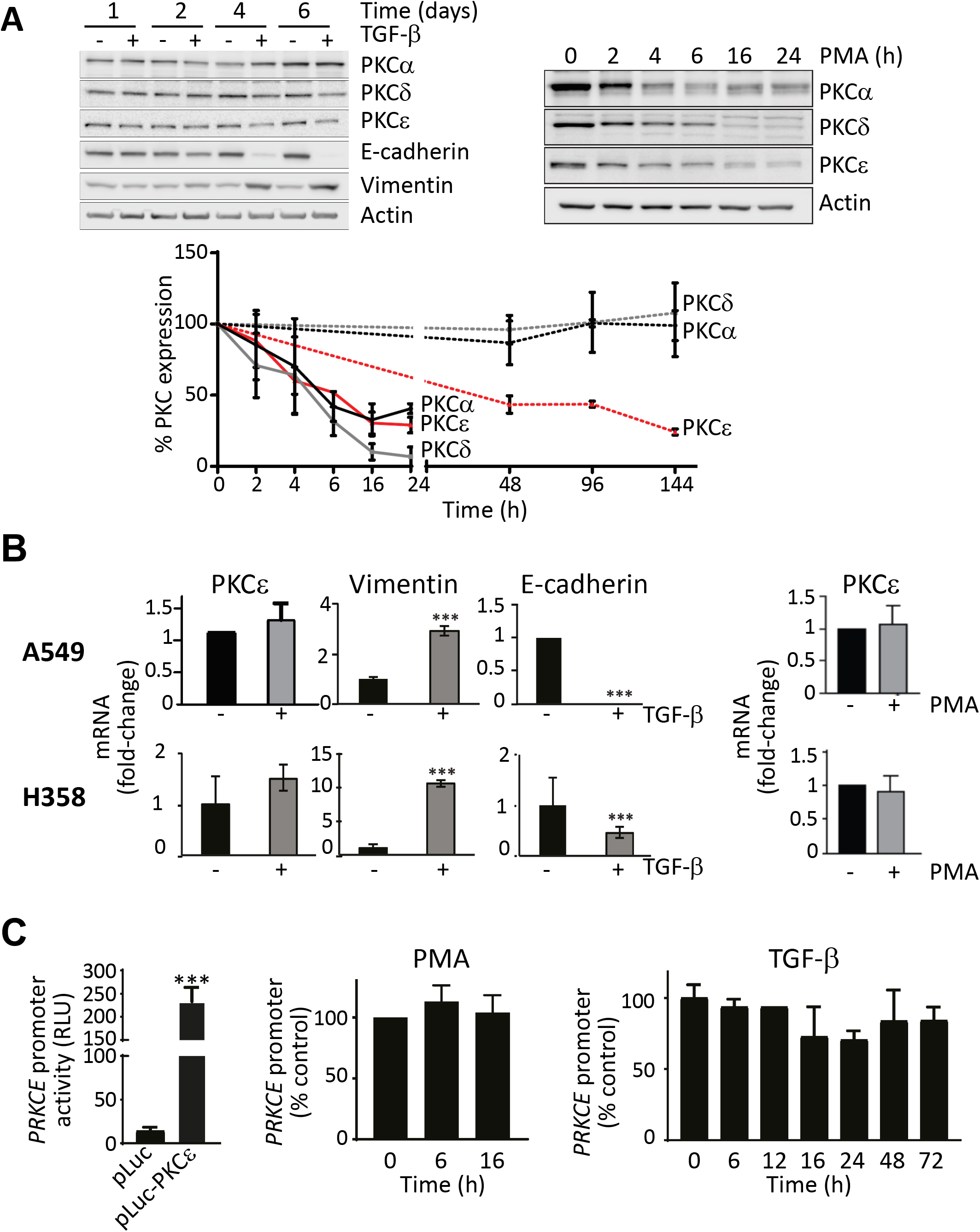
PKCε down-regulation by PMA and TGF-β in NSCLC cells. (A) Time-course analysis of PKCα, PKCδ, and PKCε protein expression in response to TGF-β (10 ng/ml, 1-6 days) or PMA (100 nM) in A549 cells. *Upper panels*, representative experiments. *Lower panel*, densitometric analysis of PKC isozyme expression, expressed as % expression relative to time = 0 h. Results are expressed as mean ± S.E.M., (3 independent experiments). (B) PKCε, vimentin and E-cadherin mRNA levels were determined by Q-PCR in A549 and H358 cells. Cells were treated with either TGF-β (10 ng/ml, 1-6 days) or PMA (100 nM, 16 h). Results are expressed as fold-change relative to untreated parental cells (mean ± S.E.M., n=3). (C) *PRKCE* promoter activity was determined in A549 cells using a reporter luciferase assay. *Left panel*, luciferase activity was determined 24 h after transfection of either empty vector or *PRKCE* luciferase vector. *Middle panel*, effect of PMA treatment. *Right panel*, effect of TGF-β (mean ± S.E.M., n=8). *RLU*, relative luciferase units.

To begin investigating the mechanisms behind PKCε down-regulation by TGF-β in NSCLC cells, we first assessed PKCε mRNA expression with Q-PCR. Using A549 and H358 NSCLC as models, we observed that PKCε mRNA levels remain essentially unchanged 6 days after TGF-β treatment. Consistent with its activity as an EMT inducer, marked E-cadherin down-regulation and a reciprocal vimentin up-regulation are detected in response to TGF-β (Fig. 1B, *left panels*). Therefore, PKCε down-regulation during TGF-β-induced mesenchymal transformation does not involve changes in PKCε mRNA synthesis or stability. In addition, no discernable changes in PKCε mRNA expression could be detected 16 h after PMA treatment in A549 or H358 cells (Fig. 1B, *right panels*), arguing that the lack of changes in PKCε mRNA levels is a general phenomenon independently of the stimuli leading to PKCε protein down-regulation.

The stable mRNA levels during PKCε down-regulation indicates that this is not due to changes at a transcriptional level. To confirm this prediction, we examined the transcriptional activity of the PKCε gene (*PRKCE*) promoter. A *PRKCE* luciferase reporter comprising bp −1933 to bp +219 bp (pLuc-PKCε) was used, since we previously demonstrated that it encompasses the relevant transcription responsive elements required for PKCε expression in cancer cells (34). Transfection of pLuc-PKCε into A549 cells resulted in a prominent reporter activity response, normalized to a *Renilla* reporter (Fig. 1C, *left panel*). Analysis of PKCε reporter activity at different times after stimulation with TGF-β (0-72 h) shows that it remained essentially unchanged. Similar results were observed in A549 cells treated with PMA for 16 h (Fig. 1C, *middle and right panel*). Altogether, these results suggest that PKCε down-regulation both during mesenchymal transformation or phorbol ester treatment does not involve changes in transcription of the *PRKCE* gene or in translational/mRNA stability mechanisms. It is therefore likely that PKCε down-regulation depends on changes in protein stability.

### Differential mechanisms involved in down-regulation of PKC isozymes in NSCLC cells

In order to begin elucidating the mechanisms involved in PKCε protein degradation, we undertook a pharmacological approach. First, we used the proteasome inhibitor bortezomib (35). We found that this agent fully prevented the down-regulation of PKCα, PKCδ and PKCε by PMA (16 h) in A549 cells (Fig. 2A). Similar studies were carried out with bafilomycin A and chloroquine, agents known to inhibit the autophagy/endolysosomal pathway (36,37). Notably, bafilomycin and chloroquine rescued the PKCε down-regulation by PMA in both A549 and H358 cells. Whereas, neither agent was capable of preventing PKCα down-regulation by the phorbol ester. A rescue of PKCδ down-regulation by chloroquine was observed only in A549 cells. The effectiveness of bafilomycin A and chloroquine in both cell lines was demonstrated by up-regulation of LC3B (Fig. 2A and 2B). A similar selective protection by bafilomycin against PMA-induced PKCε down-regulation could be observed in HCT116 colon cancer cells, SKOV3 ovarian cancer cells, PC3 prostate cancer cells and ASPC1 pancreatic cancer cells (Fig. S1). Interestingly, PKCε localization analysis revealed that PMA treatment in bafilomycin A- or chloroquine-treated A549 cells leads to the accumulation of the kinase in the periphery of large vesicular structures (Fig. 2C).

**Figure 2.**
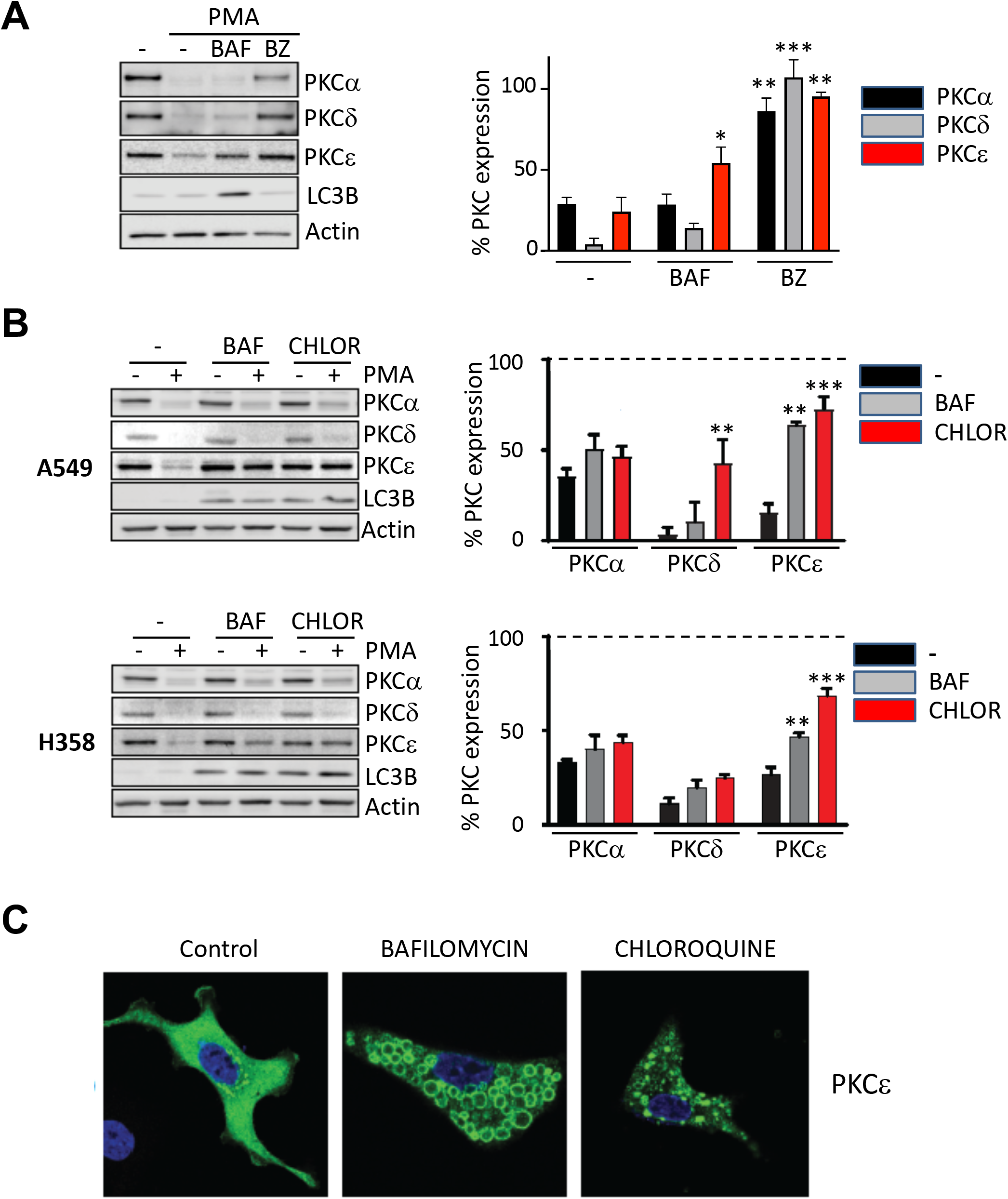
Effect of proteasome and endolysosomal inhibitors on PKC isozyme down-regulation. (A) A549 cells were serum starved for 24 h and treated with the autophagy/endolysosomal inhibitor bafilomycin A (*BAF*, 100 nM) or the proteasome inhibitor bortezomib (*BZ*, 100 nM) added 1 h before and during PMA treatment (100 nM; 16 h). (B) Autophagy/endolysosomal inhibitors bafilomycin A (*BAF*, 50 μM) or chloroquine (*CHLOR*, 50 μM) were added 1 h before PMA treatment (100 nM; 16 h). For (A) and (B), *left panel*, representative experiment; *right panel*, densitometric analysis of PKC isozyme expression, normalized by actin (mean ± S.E.M., 3 independent experiments). *Dotted line*, no PMA. *, p<0.05; **; p<0.01; ***; p<0.001 *vs*. control with no inhibitor. (C) Immunofluorescence analysis of PKCε localization. A549 cells were transfected with a FLAG-tagged PKCε expression vector (*green*) and 24 h later treated with PMA (100 nM, 16 h) in the presence or absence of either bafilomycin or chloroquine. DAPI staining was used to visualize the nucleus (*blue*). A representative experiment is shown.

When we attempted to carry out similar studies with the proteasome and autophagy/endolysosomal inhibitors to assess TGF-β-induced PKCε down-regulation, significant cell death was observed with these agents when used in incubations longer than 24 h (data not shown). Therefore, due to this toxicity we used other approaches to test the involvement of proteasome and autophagy/endolysosomal pathways during TGF-β-induced EMT (see subsequent sections). Regardless, our results using PMA strongly suggest distinctive degradation mechanisms involved in PKCε down-regulation when compared to that of PKCα and PKCδ.

### PKCε degradation is an autophagy-independent process

The protective effect of bafilomycin A and chloroquine on PKCε down-regulation suggests the potential involvement of autophagy and/or endolysosomal mechanisms. Autophagy encompasses three different processes: macroautophagy (which involves the sequestration of cytoplasmic components within double-membrane vesicles or autophagosomes, that ultimately fuses with lysosomes for degradation of its cargo), microautophagy (engulfment of cytoplasmic molecules into intralysosomal vesicles), and chaperone-mediated autophagy or CMA (which requires unfolding of the substrate for direct translocation across the lysosomal membrane) (38–40).

To address the possible contribution of the different autophagy mechanisms to PKCε degradation, we silenced key proteins of the autophagy machinery in A549 cells using RNAi. First, we knocked down specific members of the autophagy-related family (ATG) that are involved in early stages of autophagy contributing to the nucleation and elongation of autophagosomes, specifically ATG5, ATG12 and ATG6/beclin-1 (39). We observed >90% depletion of the corresponding proteins relative to non-target control (NTC), as determined by Western blot, although an unexpected cross-reactivity between ATG5 and ATG12 RNAi duplexes was also seen. Regardless, PKCε down-regulation was not affected by silencing ATG5, ATG12 or ATG6/beclin-1, neither in response to PMA (16 h) or TGF-β (48 h) treatment. The down-regulation of PKCα and PKCδ caused by PMA was also unaffected in A549 cells subjected to RNAi for these ATG proteins (Fig. 3A). A quantitative analysis of PKC levels in these experiments is depicted in Table 1A.

**Figure 3.**
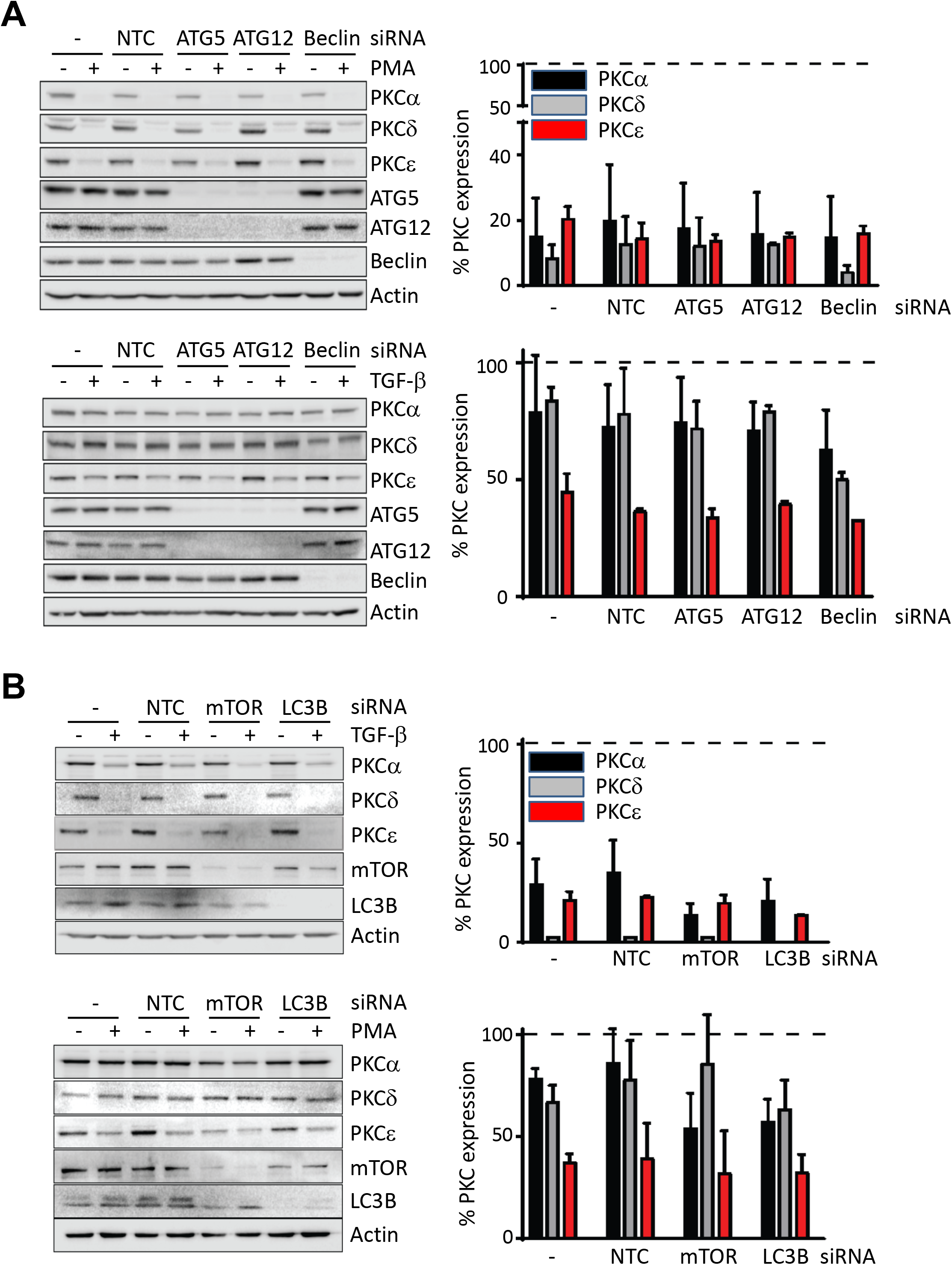
Degradation of PKC isozymes is not mediate by autophagy. A549 cells were transfected with different siRNA duplexes for autophagy proteins or non-target control (*NTC*) siRNA. Cells were serum starved for 24 h before being treated with PMA (100 nM, 16 h) or TGF-β (10 ng/ml, 48 h). (A) Effect of ATG5, ATG12 or beclin/ATG6 silencing on the expression of PKC isozymes before and after treatment with either PMA (100 nM, 16 h, *upper panel*) or TGF-β (10 ng/ml, 48 h, *lower panel*). (B) Effect of mTOR or LC3B silencing on the expression of PKC isozymes before and after treatment with either PMA (100 nM, 16 h, *upper panel*) or TGF-β (10 ng/ml, 48 h, *lower panel*). *Left panels*, representative experiments. *Right panels*, densitometric analysis of PKC isozyme expression, normalized by actin. Data are expressed as mean ± S.E.M. (n=3) relative to control cells. *Dotted line*, no PMA.

**Table 1.**
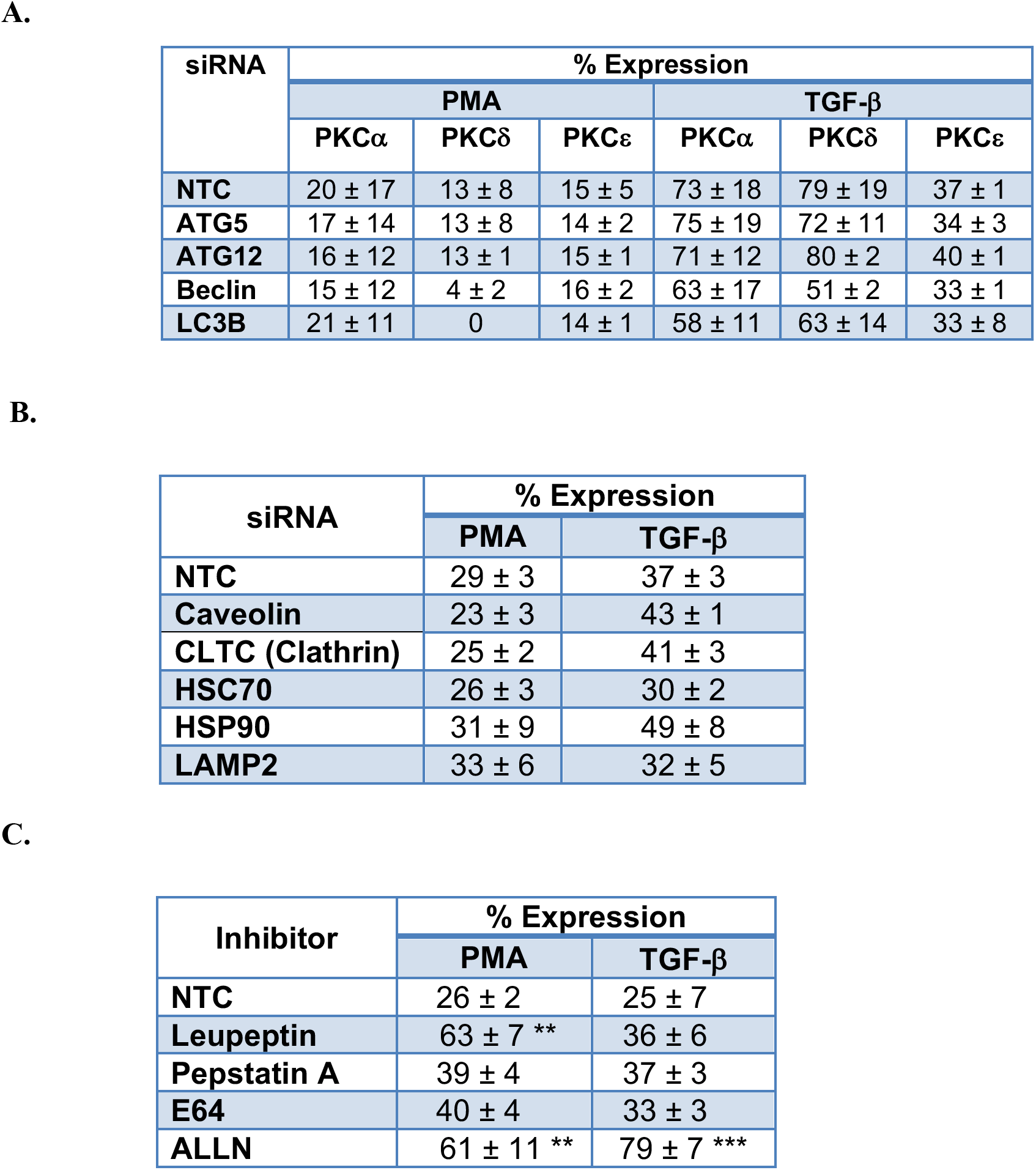
Effect of different treatments on PKC isozyme expression. (A and B) Effect of silencing different proteins on the expression of PKC isozymes after treatment with either PMA (100 nM, 16h) or TFG-β (10 ng/ml, 48 h). (B) Effect of silencing different proteins on the expression of PKCε after treatment with either PMA (100 nM, 16 h) or TFG-β (10 ng/ml, 48 h). (C) Effect of protease inhibitors on the expression of PKCε after treatment with either PMA (100 nM, 16 h) or TFG-β (10 ng/ml, 48 h). Results are expressed as % of control (untreated) cells (mean ± S.E.M., n ≥ 3 independent experiments). *NTC*, non-target control. **; p<0.01; ***; p<0.001.

| siRNA | % Expression |  |  |  |  |  |
| --- | --- | --- | --- | --- | --- | --- |
| | PMA | | | TGF-β | | |
| | PKCα | PKCδ | PKCε | PKCα | PKCδ | PKCε |
| **NTC** | 20 ± 17 | 13 ± 8 | 15 ± 5 | 73 ± 18 | 79 ± 19 | 37 ± 1 |
| **ATG5** | 17 ± 14 | 13 ± 8 | 14 ± 2 | 75 ± 19 | 72 ± 11 | 34 ± 3 |
| **ATG12** | 16 ± 12 | 13 ± 1 | 15 ± 1 | 71 ± 12 | 80 ± 2 | 40 ± 1 |
| **Beclin** | 15 ± 12 | 4 ± 2 | 16 ± 2 | 63 ± 17 | 51 ± 2 | 33 ± 1 |
| **LC3B** | 21 ± 11 | 0 | 14 ± 1 | 58 ± 11 | 63 ± 14 | 33 ± 8 |

**B.**
| siRNA | % Expression |  |
| --- | --- | --- |
| | PMA | TGF-β |
| **NTC** | 29 ± 3 | 37 ± 3 |
| **Caveolin** | 23 ± 3 | 43 ± 1 |
| **CLTC (Clathrin)** | 25 ± 2 | 41 ± 3 |
| **HSC70** | 26 ± 3 | 30 ± 2 |
| **HSP90** | 31 ± 9 | 49 ± 8 |
| **LAMP2** | 33 ± 6 | 32 ± 5 |

**C.**
| Inhibitor | % Expression |  |
| --- | --- | --- |
| | PMA | TGF-β |
| **NTC** | 26 ± 2 | 25 ± 7 |
| **Leupeptin** | 63 ± 7 ** | 36 ± 6 |
| **Pepstatin A** | 39 ± 4 | 37 ± 3 |
| **E64** | 40 ± 4 | 33 ± 3 |
| **ALLN** | 61 ± 11 ** | 79 ± 7 *** |

Next, we used RNAi to silence the expression LC3B, a protein required for the closure and fusion of autophagosomes in the late stage of autophagy (41). As observed with the early stage ATG proteins, LC3B was also dispensable for TGF-β-induced down-regulation of PKCε as well as for the down-regulation of all DAG-regulated PKCs in response to PMA. As an additional means to examine autophagy, we knocked down the mammalian target of rapamycin (mTOR), an inhibitor of autophagy (42). This approach, which should lead to stimulation of autophagy, neither led to any significant reduction in the basal levels of PKCs or modified down-regulation in response to the phorbol ester PMA or TGF-β (Fig. 3B, 4B and Table 1A).

Next, we silenced key molecules involved in CMA. First, we knocked down HSC70, a chaperone that mediates substrate targeting for CMA and also participates in endosomal microautophagy, as well as the chaperone HSP90 (38, 40). We observed that none of these proteins were required for PKCε down-regulation by either PMA or TGF-β (Fig. 4A and Table 1B). We also did not observe any changes upon knocking down LAMP-2 (lysosome-associated membrane protein 2/CD107b), the receptor in the lysosomal membrane for substrate/chaperone complexes that mediates substrate translocation into the lysosome (data not shown). This result is consistent with the fact that PKCα, PKCδ or PKCε lack the pentapeptide KFERQ-like motif required for chaperone binding (38). Altogether, these results strongly suggest that autophagy is not a mechanism involved in the degradation of PKCε or other DAG-responsive PKCs expressed in NSCLC cells.

**Figure 4.**
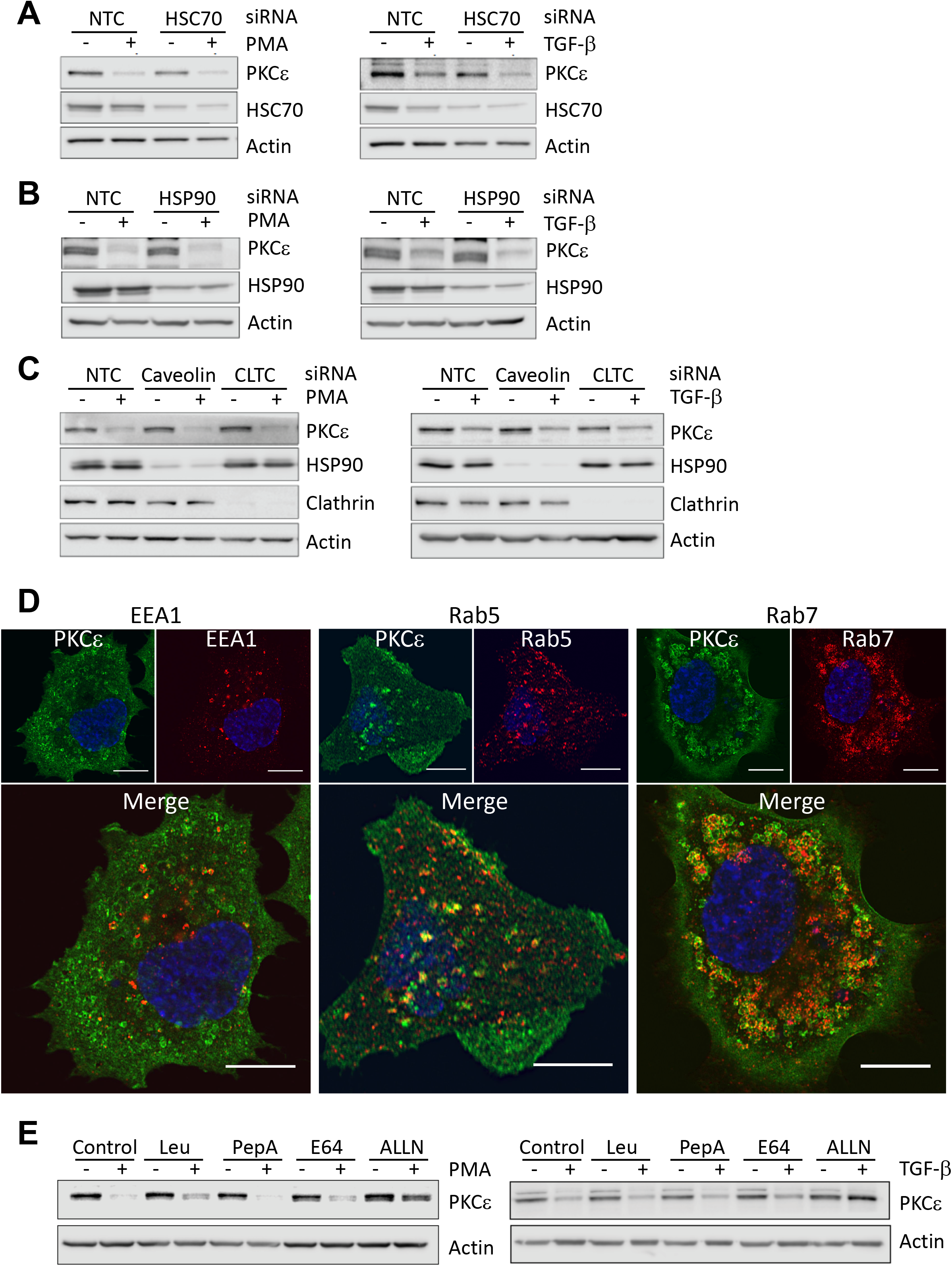
Analysis of endocytic pathways in PKCε down-regulation. (A) A549 cells were transfected with different siRNA duplexes for autophagy proteins or non-target control (*NTC*) siRNA. Cells were serum starved for 24 h before being treated with PMA (100 nM, 16 h) or TGF-β (10 ng/ml, 48 h). (A) Effect of HSC70 silencing. (B) Effect of HSP90 silencing. (C) Effect of caveolin or CLTC silencing. Representative experiments are shown. (D) A549 cells expressing FLAG-tagged PKCε were treated with PMA (100 nM, 3 h) in the presence of bafilomycin A1 (100 nM), fixed and stained for FLAG-PKCε (*green*) and endosomal markers EEA1, Rab5 and Rab7 (*red*). Nuclei was stained with DAPI (*blue*). Representative high-resolution confocal microscopy images are shown. (E) A549 cells were treated with PMA (100 nM, 16 h) or TGF-β (10 ng/ml, 48 h) in the absence or presence of leupeptin (200 μM), pepstatin A (10 μM), E64 (10 μM) or ALLN (10 μM). Expression of PKCε was determined by Western blot. A representative experiment is shown.

### PKCε degradation involves the endolysosomal pathway

Despite ruling out autophagy as a mechanism for PKCε down-regulation by silencing specific proteins of the autophagy pathways, bafilomycin A1 and chloroquine, agents widely used as inhibitors of autophagy, prevented PKCε degradation by PMA (see Fig. 2). It is known that in addition to blocking the fusion between autophagosomes and lysosomes, chloroquine also disturbs the endolysosomal machinery and bafilomycin A inhibits lysosome acidification by inhibiting the vacuolar H+ ATPase pump (V-ATPase) (36, 37). This potentially implicates an endolysosomal-dependent mechanism in PKCε down-regulation by PMA. Two main mechanisms mediate the initial steps of endocytosis that transport proteins to lysosomes, namely clathrin-mediated endocytosis (CME) and clathrin-independent endocytosis (CIE) (43, 44). To address the potential involvement of these mechanisms in PKCε down-regulation in A549 cells we silenced clathrin heavy chain 1 (CTLC/CHC1) and caveolin-1 (CAV-1), essential components of CME and CIE, respectively. Neither CTLC/CHC1 or CAV1 RNAi modified PKCε down-regulation induced by either PMA or TGF-β treatment, suggesting that CME or caveolin-dependent pathways were essentially dispensable for PKCε degradation (Fig. 4C and Table 1B). The clathrin inhibitor dynasore or the caveolin disruptor drug β-methyl cyclodextrin were also ineffective in preventing PKCε down-regulation by PMA (data not shown). These agents were toxic with incubations > 24 h and therefore could not be examined for experiments using long-term incubations with TGF-β (data not shown).

Next, to investigate the vesicular compartment(s) responsible for PKCε transport to the lysosome in response to PMA, we examined if PKCε co-localizes with specific endosomal compartment markers using confocal microscopy. Experiments were performed in the presence of bafilomycin 3 h after PMA treatment due to the significant loss of PKCε at longer times. Although under this experimental condition we detected only small co-localization with EEA1, a marker of early endosomes, we were able to observe significant co-localization with Rab5 and Rab7, markers of intermediate and late stage endocytosis, respectively. These results support the utilization of the endolysosomal pathway for the transport of PKCε to the lysosome in response to PMA (Fig. 4D). We also examined PKCε colocalization with the endosomal markers in response to TGF-β; however, we were not able to observe PKCε co-localization with endosomal markers (data not shown). It may be possible that the very slow nature of PKCε degradation under this experimental condition precludes the visualization of the association.

With the goal of identifying lysosomal protease(s) responsible for PKCε degradation we used different protease inhibitors, specifically leupeptin (for cysteine, serine and threonine peptidases), pepstatin A (for aspartyl peptidases), E64 (for cysteine peptidases) and ALLN (for calpain peptidases). Leupeptin had a preventive effect only on PKCε down-regulation by PMA, whereas E64 or pepstatin A had no significant effect on PMA- and TGF-β-induced down-regulation (Fig. 4E and Table 1C). Interestingly, ALLN rescued prevented PMA- and TGF-β-induced down-regulation. This may be related to the described inhibitory effect on proteasomal degradation (45). Altogether, these findings indicate that PKCε is degraded via two different mechanisms; the proteasome and endolysosomal pathways.

### Lysines K312 and K321 ubiquitination sites mediate PKCε degradation

As shown in Fig. 2A, treatment of A549 cells with the proteasome inhibitor bortezomib was able to prevent PKCε degradation induced by PMA. However, due to the toxicities of these agents we were unable to use them in the context of the specific degradation of PKCε induced by TGF-β. To address the potential involvement of the proteasome in PKCε degradation, we first examined if PKCε becomes ubiquitinated in response to stimulation with TGF-β. Towards this goal we co-expressed FLAG-tagged PKCε and HA-tagged ubiquitin in A549 cells, and subjected cells to treatment with TGF-β for different times (0-48 h). Upon precipitation with anti-HA antibody beads, the associated PKCε was quantified by Western blot using an anti-PKCε antibody. As shown in Fig. 5A, after 48 h of TGF-β treatment an increase in the amount of PKCε precipitating with HA-ubiquitin is observed. Therefore, TGF-β induces the ubiquitination of PKCε in A549 cells.

**Figure 5.**
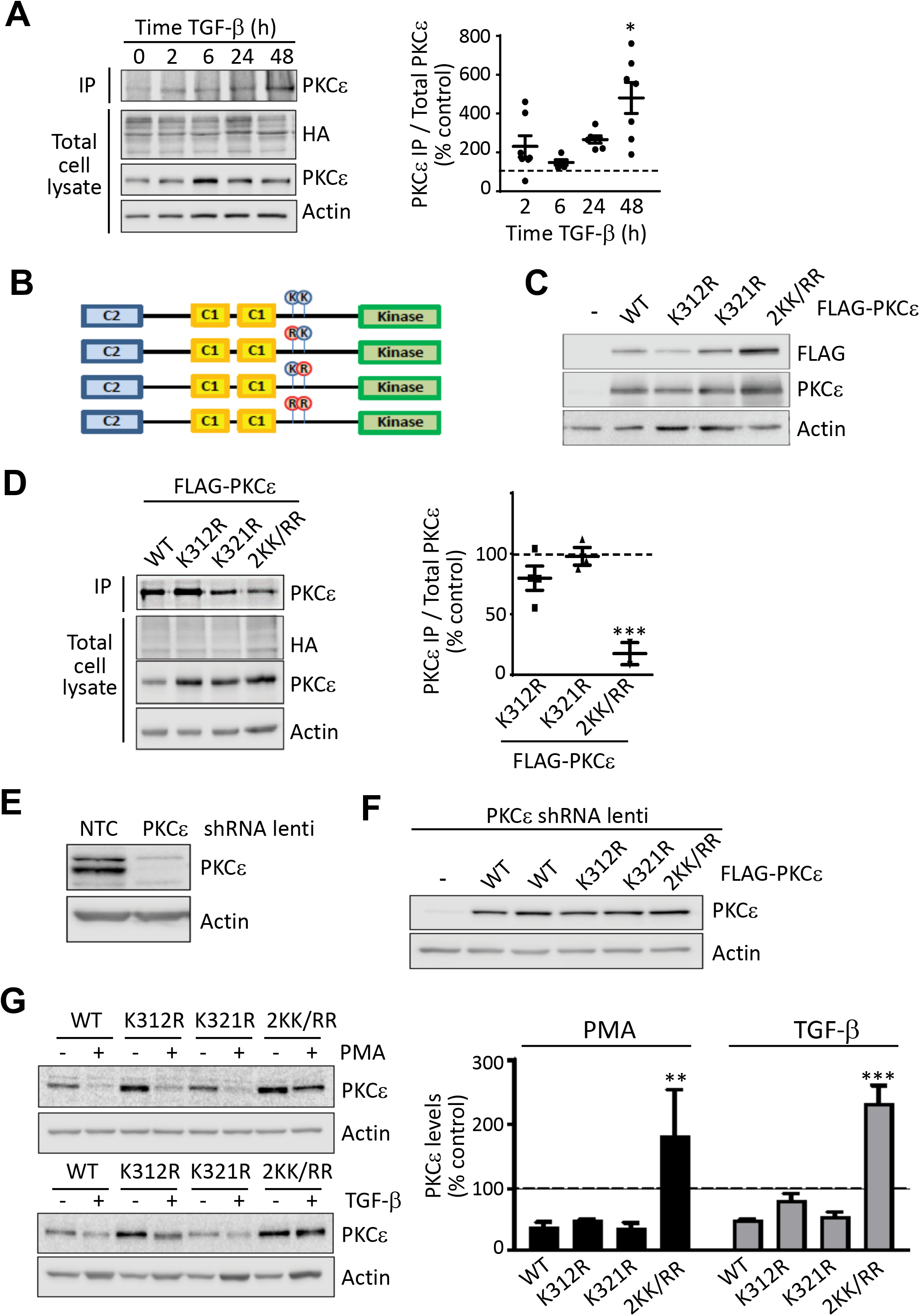
Ubiquitination is required for PKCε down-regulation. (A) Ubiquitination of FLAG-PKCε in anti-HA IPs in response to TGF-β (10 ng/ml, 0-48 h). *Left panel*, representative experiment. *Right panel*. Quantification of PKCε in IPs by densitometry, relative to total PKCε. Results were expressed as % relative to control (no TGF-β, *dotted line*) ( B) Schematic representation of PKCε mutants. (C) Expression of PKCε mutants in A549 cells by Western blot, using either anti-FLAG or anti-PKCε antibodies. (D) Ubiquitination of Flag-tagged wild-type PKCε and PKCε mutants, co-expressed with HA-tagged ubiquitin, in response to TGF-β (10 ng/ml, 48 h). *Left panel*, representative experiment. *Right panel*. Quantification of PKCε in HA IPs by densitometry, relative to total PKCε. Results were expressed as % relative to wild-type PKCε. (E) Generation of a PKCε-deficient cell line using a PKCε shRNA lentivirus. (F) Generation of A549 cell lines expressing FLAG-tagged PKCε (wild-type and mutants) in a PKCε-deficient background. (G) PKCε protein expression was analyzed by Western blot after PMA (100 nM, 16 h) or TGF-β (10 ng/ml, 48 h) treatment in A549 cell lines expressing FLAG-tagged PKCε (wild-type and mutants in a PKCε-deficient background. *Left panel*, representative experiments. *Right panel*. Quantification of PKCε by densitometry, expressed as % of control (wild-type PKCε, no treatment, *dotted line*). *, p<0.05; **; p<0.01; ***; p<0.001. *TCL*, total cell lysates; *Ip<*, immunoprecipitates.

A previous study reported that PKCε is susceptible to mono-ubiquitination at position Lys321 (46). A potential ubiquitination site in Lys312 is reported in PhosphoSitePlus® (https://www.phosphosite.org/proteinAction.action?id=1756&showAllSites=true). We generated a series of mutants to determine if these residues could be ubiquitinated in response to TGF-β. Specifically, the single mutants K321R-PKCε and K312R-PKCε, as well as the double mutant K312/K321R-PKCε (2KK/RR) (Fig. 5B). These mutants can be readily expressed in A549 cells (Fig. 5C), and in all cases they efficiently translocate to the plasma membrane in response to PMA, as determined by immunofluorescence (Fig. S2). FLAG-tagged wild-type PKCε, K312R-PKCε, K321R-PKCε and 2KK/RR-PKCε were co-expressed with HA-tagged ubiquitin in A549 cells, which were subsequently subjected to TGF-β treatment for 48 h, followed by precipitation with the anti-HA antibody. This revealed a dramatic reduction in the amount of double mutant 2KK/RR-PKCε precipitated (Fig. 5D).

Lastly, we examined the effect of mutations at Lys 312 and 321 in PKCε on TGF-β-induced degradation. In order to avoid the simultaneous expression of PKCε mutants and endogenous PKCε, we generated A549 cell lines in which endogenous PKCε was stably depleted using an shRNA lentivirus targeting the 3’ UTR (Fig. 5E). PKCε-depleted cells were then transfected with plasmids coding for FLAG tagged wild-type PKCε or the FLAG tagged PKCε mutants. Following G418 selection, stable pools for cells expressing comparable levels of wild-type PKCε, K312R-PKCε, K321R-PKCε and 2KK/RR-PKCε were generated (Fig. 5F). Cell lines were subjected to TGF-β treatment for 48 h and the expression of PKCε analyzed by Western blot. The single mutants still underwent down-regulation in response to PMA or TGF-β with a slight resistance to TGF-β induced down-regulation observed in the K312R mutant. Notably, the double mutant 2KK/RR-PKCε was resistant to down-regulation in response to PMA or TGF-β (Fig. 5G). Altogether these results suggest that TGF-β treatment induces PKCε degradation mediated by the proteasome, which depends on the ubiquitination sites Lys312 and Lys321. Ubiquitination is therefore a key mechanism involve in the degradation of PKCε triggered by TGF-β or PMA.

### Final remarks

Here, we provide evidence for the contribution of distinct mechanisms in PKC isozyme down-regulation in NSCLC cells, both in response to PMA and during the loss of expression that occurs during TGF-β induced EMT. PKCε has a pro-migratory/invasive function in cancer cells, including NSCLC cells. The pronounced down-regulation of PKCε expression during mesenchymal transformation is in some ways counterintuitive; however, it provides a permissive signal to mechanistically support the associated RhoA-mediated formation of stress fibers occurring in mesenchymally transformed NSCLC cells (31). Our studies clearly indicate that PKCε down-regulation occurring in EMT is not dependent upon transcriptional mechanisms. Indeed, while PKC isozymes have important roles in promoting the expression of key EMT transcription factors, such as Zeb1 and Twist (47–50), the inverse mechanism, *i.e.* the regulation of PKC expression by EMT transcription factors does not seem to take place, at least in the context of TGF-β-induced EMT in NSCLC cells. Rather, PKCε down-regulation in this context is mediated by proteolytic degradation of PKCε, primarily via proteasomal-dependent mechanisms.

While the ubiquitination and proteasomal-mediated degradation of different PKC family members, including PKCε, has been reported in the past (51–54), our studies provide distinctive evidence for major mechanistic differences in PKC isozyme degradation. Our initial observations revealed a selective preventative effect of chloroquine and bafilomycin on PKCε down-regulation, suggesting the existence of an additional degradation mechanism for PKCε that does not control the stability of PKCα and PKCδ, the other DAG-phorbol ester PKCs present in NSCLC cells. This may involve the utilization of endolysosomal pathways, which are sensitive to chloroquine and bafilomycin inhibition. Our studies also reveal that autophagy-mediated degradation does not seem to be involved in the down-regulation of any DAG/phorbol ester-regulated PKC, as confirmed by interfering with the expression of key components of the autophagy machinery.

## ACKNOWLEDGEMENTS

This work is supported by grants CA189765, CA196232 and ES026023 from NIH to M.G.K.

## Notes

### Competing Interest Statement

The authors have declared no competing interest.

## References

1. Griner EM, Kazanietz MG. Protein kinase C and other diacylglycerol effectors in cancer. Nat Rev Cancer. 2007;7(4):281–94.

2. Newton AC. Protein kinase C: perfectly balanced. Crit Rev Biochem Mol Biol. 2018;53(2):208–230.

3. Isakov N. Protein kinase C (PKC) isoforms in cancer, tumor promotion and tumor suppression. Semin Cancer Biol. 2018; 48:36–52.

4. Cooke M, Magimaidas A, Casado-Medrano V, Kazanietz MG. Protein kinase C in cancer: The top five unanswered questions. Mol Carcinog. 2017;56(6):1531–1542.

5. Newton AC. Regulation of the ABC kinases by phosphorylation: protein kinase C as a paradigm. Biochem J. 2003;370(Pt 2):361–71.

6. Szallasi Z, Smith CB, Pettit GR, Blumberg PM. Differential regulation of protein kinase C isozymes by bryostatin 1 and phorbol 12-myristate 13-acetate in NIH 3T3 fibroblasts. J Biol Chem. 1994;269(3):2118–24.

7. Hansra G, Garcia-Paramio P, Prevostel C, Whelan RD, Bornancin F, Parker PJ. Multisite dephosphorylation and desensitization of conventional protein kinase C isotypes. Biochem J. 1999342(Pt 2):337–44.

8. Lum MA, Barger CJ, Hsu AH, Leontieva OV, Black AR, Black JD. Protein Kinase Cα (PKCα) Is Resistant to Long Term Desensitization/Down-regulation by Prolonged Diacylglycerol Stimulation. J Biol Chem. 2016;291(12):6331–46.

9. Cooke M, Zhou X, Casado-Medrano V, Lopez-Haber C, Baker MJ, Garg R, Ann J, Lee J, Blumberg PM, Kazanietz MG. Characterization of AJH-836, a diacylglycerol-lactone with selectivity for novel PKC isozymes. J Biol Chem. 2018;293(22):8330–8341.

10. Abel EL, Angel JM, Kiguchi K, DiGiovanni J. Multi-stage chemical carcinogenesis in mouse skin: fundamentals and applications. Nat Protoc. 2009;4(9):1350–62.

11. Newton AC. Protein kinase C: perfectly balanced. Crit Rev Biochem Mol Biol. 2018;53(2):208–230.

12. Garg R, Benedetti LG, Abera MB, Wang H, Abba M, Kazanietz MG. Protein kinase C and cancer: what we know and what we do not. Oncogene. 2014;33(45):5225–37.

13. Newton AC. Protein kinase C: perfectly balanced. Crit Rev Biochem Mol Biol. 2018;53(2):208–230.

14. Gobbi G, Mirandola P, Sponzilli I, Micheloni C, Malinverno C, Cocco L, Vitale M. Timing and expression level of protein kinase C epsilon regulate the megakaryocytic differentiation of human CD34 cells. Stem Cells. 2007;25(9):2322–9.

15. Denning MF, Dlugosz AA, Williams EK, Szallasi Z, Blumberg PM, Yuspa SH. Specific protein kinase C isozymes mediate the induction of keratinocyte differentiation markers by calcium. Cell Growth Differ. 1995;6(2):149–57.

16. Jain K, Basu A. The Multifunctional Protein Kinase C-ε in Cancer Development and Progression. Cancers (Basel). 2014;6(2):860–78.

17. Gorin MA, Pan Q. Protein kinase C epsilon: an oncogene and emerging tumor biomarker. Mol Cancer. 2009 Feb 19;8:9.

18. Basu A, Sivaprasad U. Protein kinase Cepsilon makes the life and death decision. Cell Signal. 2007;19(8):1633–42.

19. Garg R, Blando JM, Perez CJ, Abba MC, Benavides F, Kazanietz MG. Protein Kinase C Epsilon Cooperates with PTEN Loss for Prostate Tumorigenesis through the CXCL13-CXCR5 Pathway. Cell Rep. 2017;19(2):375–388.

20. Garg R, Blando J, Perez CJ, Wang H, Benavides FJ, Kazanietz MG. Activation of nuclear factor κB (NF-κB) in prostate cancer is mediated by protein kinase C epsilon (PKCepsilon). J Biol Chem. 2012;287(44):37570–82.

21. Caino MC, Lopez-Haber C, Kissil JL, Kazanietz MG. Non-small cell lung carcinoma cell motility, rac activation and metastatic dissemination are mediated by protein kinase C epsilon. PLoS One. 2012;7(2):e31714.

22. Aziz MH, Manoharan HT, Church DR, Dreckschmidt NE, Zhong W, Oberley TD, Wilding G, Verma AK. Protein kinase Cepsilon interacts with signal transducers and activators of transcription 3 (Stat3), phosphorylates Stat3Ser727, and regulates its constitutive activation in prostate cancer. Cancer Res. 2007;67(18):8828–38.

23. Garg R, Blando JM, Perez CJ, Lal P, Feldman MD, Smyth EM, Ricciotti E, Grosser T, Benavides F, Kazanietz MG. COX-2 mediates pro-tumorigenic effects of PKCε in prostate cancer. Oncogene. 2018;37(34):4735–4749.

24. Bae KM, Wang H, Jiang G, Chen MG, Lu L, Xiao L. Protein kinase C epsilon is overexpressed in primary human non-small cell lung cancers and functionally required for proliferation of non-small cell lung cancer cells in a p21/Cip1-dependent manner. Cancer Res. 2007;67(13):6053–63.

25. Caino MC, Lopez-Haber C, Kim J, Mochly-Rosen D, Kazanietz MG. Proteins kinase Cɛ is required for non-small cell lung carcinoma growth and regulates the expression of apoptotic genes. Oncogene. 2012;31(20):2593–600.

26. Bao L, Gorin MA, Zhang M, Ventura AC, Pomerantz WC, Merajver SD, Teknos TN, Mapp AK, Pan Q. Preclinical development of a bifunctional cancer cell homing, PKCepsilon inhibitory peptide for the treatment of head and neck cancer. Cancer Res. 2009;69(14):5829–34.

27. Hafeez BB, Zhong W, Weichert J, Dreckschmidt NE, Jamal MS, Verma AK. Genetic ablation of PKC epsilon inhibits prostate cancer development and metastasis in transgenic mouse model of prostate adenocarcinoma. Cancer Res. 2011;71(6):2318–27.

28. Dann SG, Golas J, Miranda M, Shi C, Wu J, Jin G, Rosfjord E, Upeslacis E, Klippel A. p120 catenin is a key effector of a Ras-PKCɛ oncogenic signaling axis. Oncogene. 2014;33(11):1385–94.

29. Garg R, Cooke M, Benavides F, Abba MC, Cicchini M, Feldser DM, Kazanietz MG. PKC epsilon is required for KRAS-driven lung tumorigenesis. Cancer Res. 2020 Sep 29:canres.1300.2020. doi: 10.1158/0008-5472.CAN-20-1300. Epub ahead of print.

30. Jain K, Basu A. Protein Kinase C-ε Promotes EMT in Breast Cancer. Breast Cancer (Auckl). 2014;8:61–7.

31. Casado-Medrano V, Barrio-Real L, Wang A, Cooke M, Lopez-Haber C, Kazanietz MG. Distinctive requirement of PKCε in the control of Rho GTPases in epithelial and mesenchymally transformed lung cancer cells. Oncogene. 201938(27):5396–5412.

32. Lopez-Haber C, Barrio-Real L, Casado-Medrano V, Kazanietz MG. Heregulin/ErbB3 Signaling Enhances CXCR4-Driven Rac1 Activation and Breast Cancer Cell Motility via Hypoxia-Inducible Factor 1α. Mol Cell Biol. 2016;36(15):2011–26.

33. Barrio-Real L, Lopez-Haber C, Casado-Medrano V, Goglia AG, Toettcher JE, Caloca MJ, Kazanietz MG. P-Rex1 is dispensable for Erk activation and mitogenesis in breast cancer. Oncotarget. 2018;9(47):28612–28624.

34. Wang H, Gutierrez-Uzquiza A, Garg R, Barrio-Real L, Abera MB, Lopez-Haber C, Rosemblit C, Lu H, Abba M, Kazanietz MG. Transcriptional regulation of oncogenic protein kinase Cϵ (PKCϵ) by STAT1 and Sp1 proteins. J Biol Chem. 2014;289(28):19823–38.

35. Manasanch EE, Orlowski RZ. Proteasome inhibitors in cancer therapy. Nat Rev Clin Oncol. 2017;14(7):417–433.

36. Mauvezin C, Neufeld TP. Bafilomycin A1 disrupts autophagic flux by inhibiting both V-ATPase-dependent acidification and Ca-P60A/SERCA-dependent autophagosome-lysosome fusion. Autophagy. 2015;11(8):1437–8.

37. Pasquier B. Autophagy inhibitors. Cell Mol Life Sci. 2016;73(5):985–1001.

38. Yim WW, Mizushima N. Lysosome biology in autophagy. Cell Discov. 2020 Feb 11;6:6.

39. Lamb CA, Yoshimori T, Tooze SA. The autophagosome: origins unknown, biogenesis complex. Nat Rev Mol Cell Biol. 2013;14(12):759–74.

40. Levy JMM, Towers CG, Thorburn A. Targeting autophagy in cancer. Nat Rev Cancer. 2017;17(9):528–542.

41. Schaaf MB, Keulers TG, Vooijs MA, Rouschop KM. LC3/GABARAP family proteins: autophagy-(un)related functions. FASEB J. 2016;30(12):3961–3978.

42. Janku F, McConkey DJ, Hong DS, Kurzrock R. Autophagy as a target for anticancer therapy. Nat Rev Clin Oncol. 2011;8(9):528–39.

43. Kaksonen M, Roux A. Mechanisms of clathrin-mediated endocytosis. Nat Rev Mol Cell Biol. 2018;19(5):313–326.

44. Sandvig K, Kavaliauskiene S, Skotland T. Clathrin-independent endocytosis: an increasing degree of complexity. Histochem Cell Biol. 2018;150(2):107–118.

45. Yeung SJ, Chen SH, Chan L. Ubiquitin-proteasome pathway mediates intracellular degradation of apolipoprotein B. Biochemistry. 1996;35(43):13843–8.

46. Yang W, Xia Y, Cao Y, Zheng Y, Bu W, Zhang L, You MJ, Koh MY, Cote G, Aldape K, Li Y, Verma IM, Chiao PJ, Lu Z. EGFR-induced and PKCε monoubiquitylation-dependent NF-κB activation upregulates PKM2 expression and promotes tumorigenesis. Mol Cell. 2012;48(5):771–84.

47. Tam WL, Lu H, Buikhuisen J, Soh BS, Lim E, Reinhardt F, Wu ZJ, Krall JA, Bierie B, Guo W, Chen X, Liu XS, Brown M, Lim B, Weinberg RA. Protein kinase C α is a central signaling node and therapeutic target for breast cancer stem cells. Cancer Cell. 2013;24(3):347–64.

48. Llorens MC, Rossi FA, García IA, Cooke M, Abba MC, Lopez-Haber C, Barrio-Real L, Vaglienti MV, Rossi M, Bocco JL, Kazanietz MG, Soria G. PKCα Modulates Epithelial-to-Mesenchymal Transition and Invasiveness of Breast Cancer Cells Through ZEB1. Front Oncol. 2019;9:1323.

49. Tedja R, Roberts CM, Alvero AB, Cardenas C, Yang-Hartwich Y, Spadinger S, Pitruzzello M, Yin G, Glackin CA, Mor G. Protein kinase Cα-mediated phosphorylation of Twist1 at Ser-144 prevents Twist1 ubiquitination and stabilizes it. J Biol Chem. 2019;294(13):5082–5093.

50. Rahimova N, Cooke M, Zhang S, Baker MJ, Kazanietz MG. The PKC universe keeps expanding: From cancer initiation to metastasis. Adv Biol Regul. 2020;78:100755. doi: 10.1016/j.jbior.2020.100755. Epub ahead of print.

51. Lee HW, Smith L, Pettit GR, Vinitsky A, Smith JB. Ubiquitination of protein kinase C-alpha and degradation by the proteasome. J Biol Chem. 1996;271(35):20973–6.

52. Lee HW, Smith L, Pettit GR, Smith JB. Bryostatin 1 and phorbol ester down-modulate protein kinase C-alpha and -epsilon via the ubiquitin/proteasome pathway in human fibroblasts. Mol Pharmacol. 1997;51(3):439–47.

53. Lu Z, Liu D, Hornia A, Devonish W, Pagano M, Foster DA. Activation of protein kinase C triggers its ubiquitination and degradation. Mol Cell Biol. 1998;18(2):839–45.

54. Leontieva OV, Black JD. Identification of two distinct pathways of protein kinase Calpha down-regulation in intestinal epithelial cells. J Biol Chem. 2004;279(7):5788–801.

