## Supplemental Figures for "Mechanisms of protein kinase C epsilon down-regulation by transforming growth factor-beta in lung cancer cells"

### SUPPLEMENTAL FILES

### SUPPLEMENTARY FIGURES

**Figure S1. Protection of PKCε down-regulation in different cell lines.** The indicated cell lines were serum starved for 24 h and treated with bafilomycin A (*BAF*, 100 nM) added 1 h before and for the duration of PMA treatment (100 nM, 16 h). A representative experiment for the expression of PKC isozymes is shown.

**Figure S2. Translocation of FLAG-tagged PKCε (wild-type and mutants) in response to PMA.** Immunofluorescence analysis of PKCε mutant localization in A549 cells in response to PMA (100 nM, 30min), using an anti-FLAG antibody (*green*). Nuclei were stained with DAPI (*blue*). A representative experiment is shown.

**Figure S1**

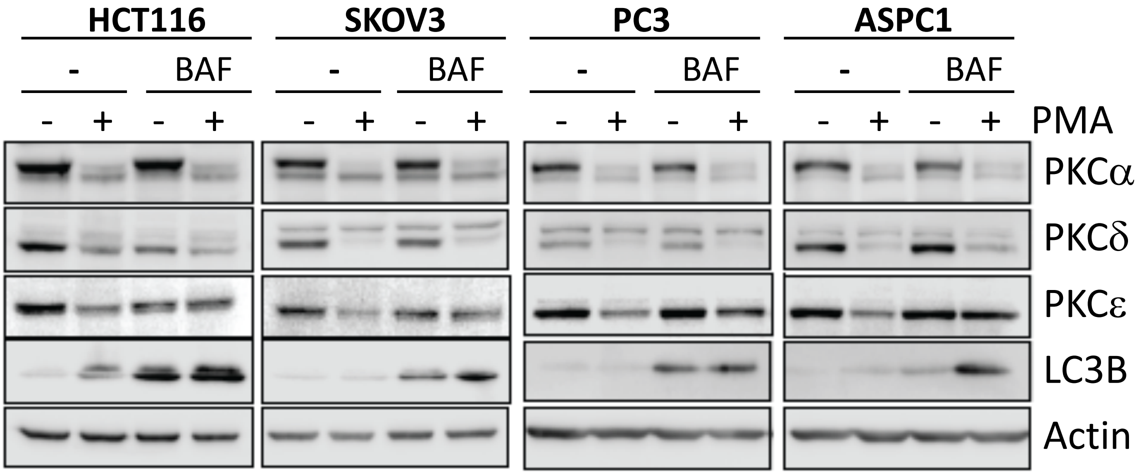

HCT116
SKOV3
PC3
ASPC1
-
BAF
-
BAF
-
BAF
-
BAF
- + - + - + - + - + - + - + - + PMA
PKCα
PKCδ
PKCε
LC3B
Actin

**Figure S2**

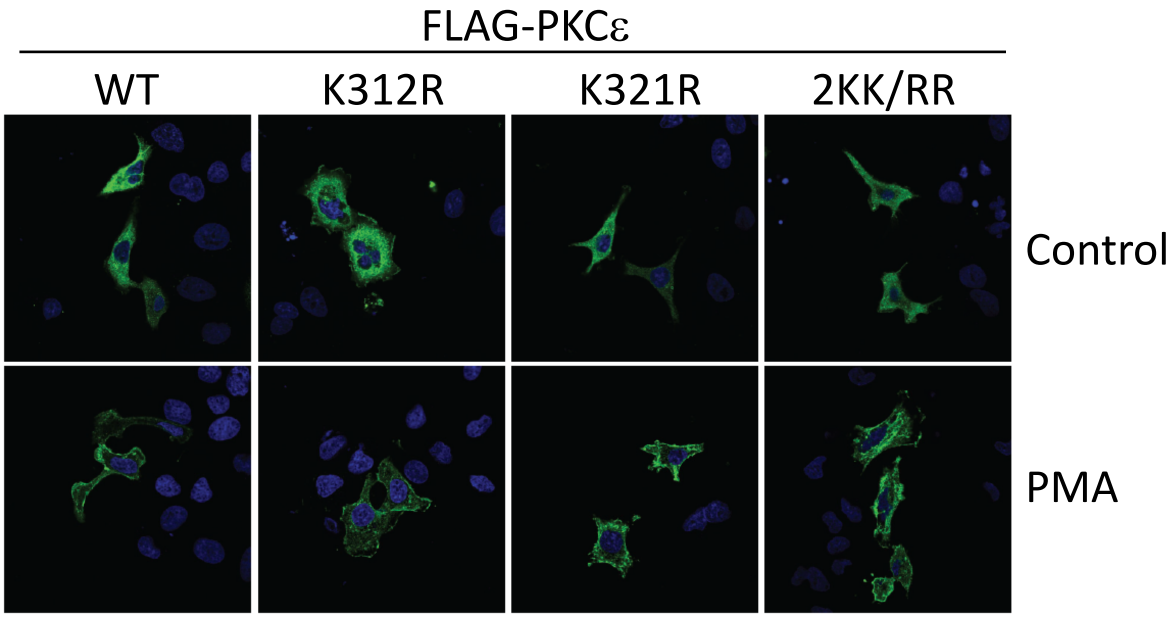
